# Inflammatory macrophages prevent colonic goblet and enteroendocrine cell differentiation through Notch signaling

**DOI:** 10.1101/2023.06.29.547119

**Authors:** Roger Atanga, Aaron S. Romero, Anthony Jimenez Hernandez, Eduardo Peralta-Herrera, Seth D. Merkley, Julie G. In, Eliseo F. Castillo

## Abstract

Inflammatory macrophages in the intestine are a key pathogenic factor driving inflammatory bowel disease (IBD). Here, we report the role of inflammatory macrophage-mediated notch signaling on secretory lineage differentiation in the intestinal epithelium. Utilizing IL-10-deficient (*Il10^-/-^*) mice, a model of spontaneous colitis, we found an increase in Notch activity in the colonic epithelium as well as an increase in intestinal macrophages expressing Notch ligands, which are increased in macrophages upon inflammatory stimuli. Furthermore, a co-culture system of inflammatory macrophages and intestinal stem and proliferative cells during differentiation reduced goblet and enteroendocrine cells. This was recapitulated when utilizing a Notch agonist on human colonic organoids (colonoids). In summary, our findings indicate that inflammatory macrophages upregulate notch ligands that activate notch signaling in ISC via cell-cell interactions, which in turn inhibits secretory lineage differentiation in the gastrointestinal (GI) tract.

## Introduction

Notch signaling is a cell-cell communication mechanism important for many aspects of biology, including intestinal stem cell (ISC) differentiation^1,2^. This process mainly occurs through signal-sending cells expressing Notch ligands that bind Notch receptors on signal-receiving cells. There are 5 mammalian Notch ligands (Delta-like: DLL1, DLL3, and DLL4; and Serrate (Jagged)-like: Jagged1 and Jagged2) and 4 Notch receptors (Notch1–4)^2^. Notch activation occurs after i) Notch ligand binds a Notch receptor resulting in proteolysis of the extracellular domain of Notch receptor by the metalloprotease ADAM10^3^ and ii) the mechanical pull of Notch ligand on the Notch receptor. The Notch intracellular domain (NICD) is then released into the cytosol by the γ-secretase complex. The cytosolic NICD translocate to the nucleus binding the transcription factor CSL (CBF-1 /Suppressor of Hairless/ Lag-1) resulting in the expression of numerous target genes such as *Hes1* (Hairy and enhancer of split 1) and *Hey1* (Hes related family bHLH transcription factor with YRPW motif 1) that mediate Notch activity^1,2,4^. During ISC differentiation, continuous Notch signaling through *Hes1* and *Hey1* expression represses the transcription factor *Atoh1* (atonal bHLH transcription factor 1) to promote absorptive cellular differentiation, *e.g*., enterocytes in small intestine and colonocytes in the colon ^4-6^. Whereas, certain secretory cells, like goblet and enteroendocrine cells, are promoted through the inhibition of Notch signaling and the expression of *Atoh1* ^7-9^. Nevertheless, both absorptive and secretory cells are critical to the epithelial barrier to maintain intestinal homeostasis.

Intestinal macrophages also contribute to ISC renewal, homeostasis, and differentiation, as depletion of macrophages reduces stem cell numbers and impairs cellular differentiation ^10^. Macrophages are abundant in the intestine and act as a first line of defense in the gut. Unlike other tissue macrophages, intestinal macrophages exhibit an anti-inflammatory phenotype and display high phagocytic and bactericidal activity, as well as a high tolerance towards foreign material ^11,12^. However, in states of chronic intestinal inflammation, like Crohn’s disease (CD) and ulcerative colitis (UC) the two major forms of inflammatory bowel disease (IBD), an accumulation of inflammatory macrophages can be observed in the mucosa ^13,14^. The influence of inflammatory macrophages on ISC differentiation is unclear but macrophages have been found to be both Notch signal-sending and signal-receiving cells. Interestingly, Notch signaling is dysregulated in IBD^5,15-18^. Nevertheless, it is unclear which cell-type promotes increased Notch signaling in IBD. Herein, we show intestinal macrophages in the inflamed colon of a mouse model of IBD display increased expression of Notch ligands Jagged1 and DLL3. The inflamed colon also showed increased levels of Notch target gene expression and reduced levels of *Atoh1*. We further show inflammatory macrophages prevent secretory cell differentiation while anti-inflammatory macrophages promoted goblet and enteroendocrine cell differentiation. This was recapitulated in human organoids in the absence of macrophages and through the utilization of a Notch agonist. In summary, our findings indicate that inflammatory macrophages can inhibit secretory lineage differentiation in the gastrointestinal (GI) tract through Notch signaling.

## Results and Discussion

### Increased Notch activity in the inflamed colonic epithelium

Notch signaling is a major pathway dictating absorptive and secretory cellular differentiation. High Notch activity is observed in the crypts of the inflamed colon of UC patients^18,19^. UC patients also show decreased *ATOH1* expression as well as decreased goblet cell numbers and mucus^5^. CD also shows a similar loss of secretory cells^17^. In mice, the loss of Notch signaling via genetic or chemical inhibition results in intestinal inflammation^1^. Similar to IBD, *Il10*^-/-^ mice also have reduced goblet cells and mucus^20^. The *Il10*^-/-^ mouse model is representative of chronic, immune-mediated intestinal inflammation as IL-10 polymorphisms confer increased risk to both CD and UC ^21,22^. We were interested to see if Notch target genes were upregulated in the inflamed colon of *Il10*^-/-^ mice. Utilizing fecal lipocalin-2 (LCN-2) as a marker of intestinal inflammation^23-26^, we observed increased fecal LCN-2 levels in *Il10^-/-^* mice when compared to control C57BL/6 (B6) mice (**Suppl. Fig. 1 A**) as well as increased IFNγ, IL-17A, and IL-22 production from CD4^+^ T cells isolated from mesenteric lymph nodes (**Suppl. Fig. 1 D-F**). Next, we isolated colonocytes from the inflamed colon of *Il10*^-/-^ mice and observed an increase in several Notch target genes including *Hes1*, *Hey1*, and *Dtx1* when compared to the uninflamed colon of control B6 mice (**Fig. 1 A-C**). Although not significantly different from controls, the Notch target gene *Hes5* was trending upwards (**Fig. 1 D**). Examination of *Gli1*, a gene associated with Hedgehog signaling, another cell determination pathway, showed no change in expression (**Fig. 1 E**). However, when we examined *Atoh1*, a promoter of secretory cells, colonocytes from *Il10*^-/-^ mice showed a significant decrease in *Atoh1* expression compared to the uninflamed colon (**Fig. 1 F**). To determine if this change in Notch activity is due to lack of IL-10 or chronic inflammation, we compared *Hes1* and *Atoh1* expression in an acute and chronic model of dextran sodium sulfate (DSS) induced colitis. Isolated colonocytes from chronic DSS B6 mice showed similar expression patterns of both *Hes1* and *Atoh1* to colitic *Il10*^-/-^ mice (**Suppl. Fig. 1 B, C**). However, colonic *Hes1* and *Atoh1* expression in the acute DSS model were similar to untreated B6 mice (**Suppl. Fig. 1 B, C**). Our data supports prior evidence^5,15-18^ of enhanced Notch activity in the colonic epithelium during chronic intestinal inflammation.

**Figure 1.**
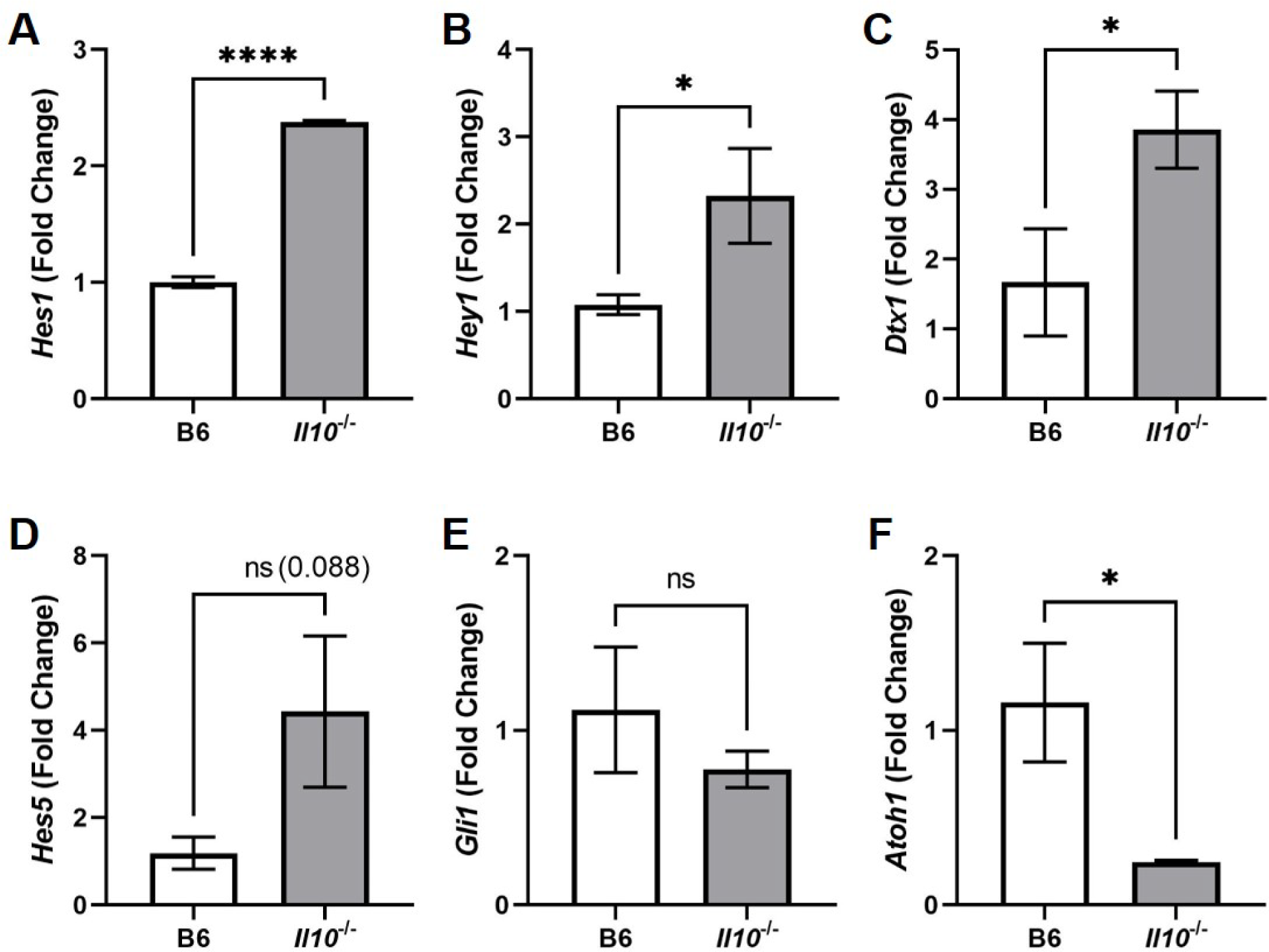
Notch target gene expression in the colon of *Il10*^-/-^ mice. mRNA expression of **(A)** *Hes1*, **(B)** *Hey1*, **(C)** *Dtx1*, **(D)** *Hes5*, **(E)** *Gli1*, and **(F)** *Atoh1* in the inflamed colon of *Il10*^-/-^ mice compared to the uninflamed colon of (B6) mice. Two independent experiments, n = 8/group. Graphs indicate mean (±SEM). * P<0.05, ****P<0.0001, ns, not significant. Significance was calculated by two-tailed unpaired Student’s t test.

### Colonic macrophages in the inflamed colon express enhanced levels of Notch ligands

Amassing evidence from IBD patient samples, animal and mathematical models of IBD, as well as identified IBD susceptible risk loci, have all suggested that macrophages are a major cell-type contributing to IBD pathogenesis^5,12-18,27-30^. Given the abundance of macrophages in the gut, we next sought to identify features of colonic macrophages that could contribute to the observed Notch activity in the inflamed colon. As we have previously reported^26^, we isolated colonic macrophages from the lamina propria (LP). Similar to what has been observed in the mucosa of IBD patients^13,14^, there was an overall increase in the total number of LP macrophages (CX3CR1^+^MHCII^+^CD68^+^) in *Il10*^-/-^ mice compared to controls (**Suppl. Fig. 2 A-C**). A significant portion of these LP macrophages from colitic *Il10*^-/-^ mice expressed Jagged1 compared to macrophages in the non-inflamed gut (**Fig. 2 A, B**). There was also an increase in DLL3^+^ LP macrophages in *Il10*^-/-^ mice (**Fig. 2 C, D**). However, Jagged2, DLL1 and DLL4 expression in macrophages did not show a significant difference between both groups of mice (**Suppl. Fig. 2 D-F**). Lastly, this population of macrophages isolated from colitic mice also showed increased levels of cell surface TLR4 expression (**Suppl. Fig. 2 A, B**) which has been reported to be upregulated during intestinal inflammation^31^.

**Figure 2.**
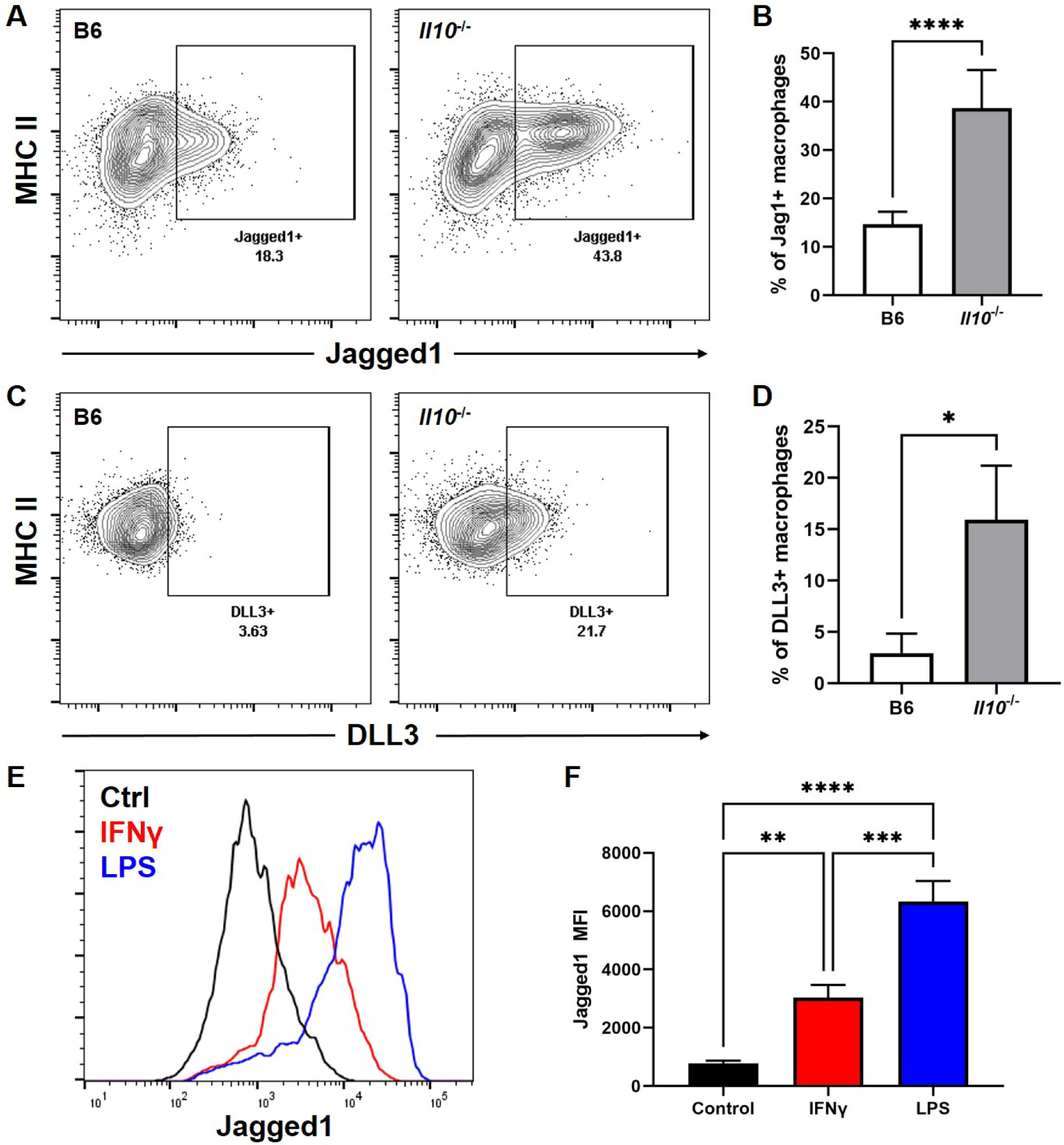
Notch ligand expression in lamina propria macrophages. Phenotype of lamina propria macrophages in colitic *Il10*^-/-^ mice**. (A)** Representative flow cytometric plots showing MHC II and Jagged1 expression in LP macrophages (gated on CD45^+^MHC-II^+^CX3CR1^+^CD68^+^ cells) isolated from colitic *Il10*^-/-^ mice and non-colitic B6 mice. **(B)** Graph showing percent of Jagged1^+^ (Jag1) LP macrophages from both group of mice. **(C)** Representative flow cytometric plots showing MHC II and DLL3 expression in LP macrophages isolated from colitic *Il10*^-/-^ mice and non-colitic B6 mice. **(D)** Graph showing percent of DLL3^+^ LP macrophages from both group of mice. **(E)** Representative histogram showing Jagged1 expression in B6 macrophages stimulated with LPS (Blue) or IFNγ (Red) compared to baseline/unstimulated (Black) macrophages. **(F)** Graph showing the geometric mean fluorescence intensity (MFI) of Jagged1 on unstimulated and stimulated B6 macrophages. Representative of two independent experiments, Graphs indicate mean (±SD). * P<0.05, **P<0.01, ***P<0.001, ****P<0.0001. Significance was calculated by two-tailed unpaired Student’s t test or one-way ANOVA.

Upon observing that inflammatory macrophages from *Il10*^-/-^ mice show increased Notch ligand expression *in vivo*, we wanted to determine if inflammatory stimuli could upregulate Notch ligand expression in macrophages. Therefore, we generated macrophages from bone marrow monocytes isolated from B6 mice as we have previously described ^26,32,33^. Macrophages were stimulated overnight with lipopolysaccharide (LPS) or interferon-gamma (IFNγ) to examine changes in Notch ligand expression. Flow cytometric analysis revealed both LPS and IFNγ stimuli increased expression of Jagged1 **(Fig. 2 E, F).** Under our conditions, these stimuli also increased Jagged2, DLL3 and DLL4 but did not alter DLL1 expression compared to unstimulated macrophages **(Suppl. Fig. 3 A-D)**. Taken together, our data demonstrates an inflammatory microenvironment and signals increase Notch ligand expression in macrophages.

### Inflammatory macrophages prevent secretory lineage differentiation

At steady state, intestinal macrophages contribute to ISC differentiation^10^; however, it is unclear how inflammatory macrophages affect ISC differentiation. We reveal a compelling correlation of increased colonic epithelial Notch activity and Notch ligand expression by colonic macrophages in the inflamed colon of *Il10*^-/-^ mice. One possibility is that the inflamed colon harbor ISC that have enhanced expression of Notch receptors that increases Notch activity. When colonic Lgr5^+^ crypt cells were examined for Notch receptor expression there were no observed differences between the proportion of cells expressing Notch receptors (Notch1-4) in the inflamed and uninflamed crypt (**Suppl. Fig. 4 A-E**). This suggest an increase in Notch ligand expression in Notch signal-sending cell(s) may increase Notch activity in the inflamed colon. Interestingly, Notch-1 and -2 as well as DLL1 and DLL4 have been reported to be the respective key receptors and ligands regulating intestinal homeostasis^34-37^. Given there was no difference in DLL1 and DLL4 expression in macrophages suggest other Notch ligand-positive cells contribute to Notch signaling during homeostatic conditions.

We next wanted to examine the impact of macrophage-mediated Notch signaling on colonic ISC differentiation. To do so, we set-up a co-culture system^38^ (**Fig. 3**) allowing colonic intestinal stem cells to differentiate in the presence of anti-inflammatory or inflammatory macrophages. Transwells were seeded with ISC derived from B6 colonoids to form a monolayer. After reaching confluency, monolayers were cultured with either anti-inflammatory (unstimulated, **Fig. 3 A4, C4**) or inflammatory (IFNγ stimulated, **Fig. 3 A5, C5**) macrophages on the basolateral side of the transwell. As shown in **Fig. 2 E** and **Suppl. Fig. 3A-D**, inflammatory macrophages express high levels of Notch ligands. ISC monolayers co-cultured with anti-inflammatory macrophages showed a significant increase in goblet cells as stained by Muc2 (**Fig. 3 A2**) and enteroendocrine cells marked by chromogranin A (CHGA) (**Fig. 3 C2**) when compared to control monolayers with no macrophages (**Fig. 3 A1, C1**). On the other hand, monolayers cultured with inflammatory macrophages inhibited goblet (**Fig. 3 A3, B**) and enteroendocrine cell (**Fig. 3 C3, D**) differentiation. Upon quantification, both goblet and enteroendocrine cells increased by two folds upon co-culture with anti-inflammatory macrophages while there was a slight decrease in goblet cells and no change in enteroendocrine cells upon co-culture with inflammatory macrophages **(Fig. 3 B, D)**. The effect of inflammatory macrophages on the barrier appeared surprisingly similar to monocultures; however, monolayers cultured with inflammatory macrophages showed changes in phalloidin (**Fig. 3 A3, C3**) staining. Although not the scope of this study, in the future we will examine how inflammatory macrophages impact epithelial maturation.

**Figure 3.**
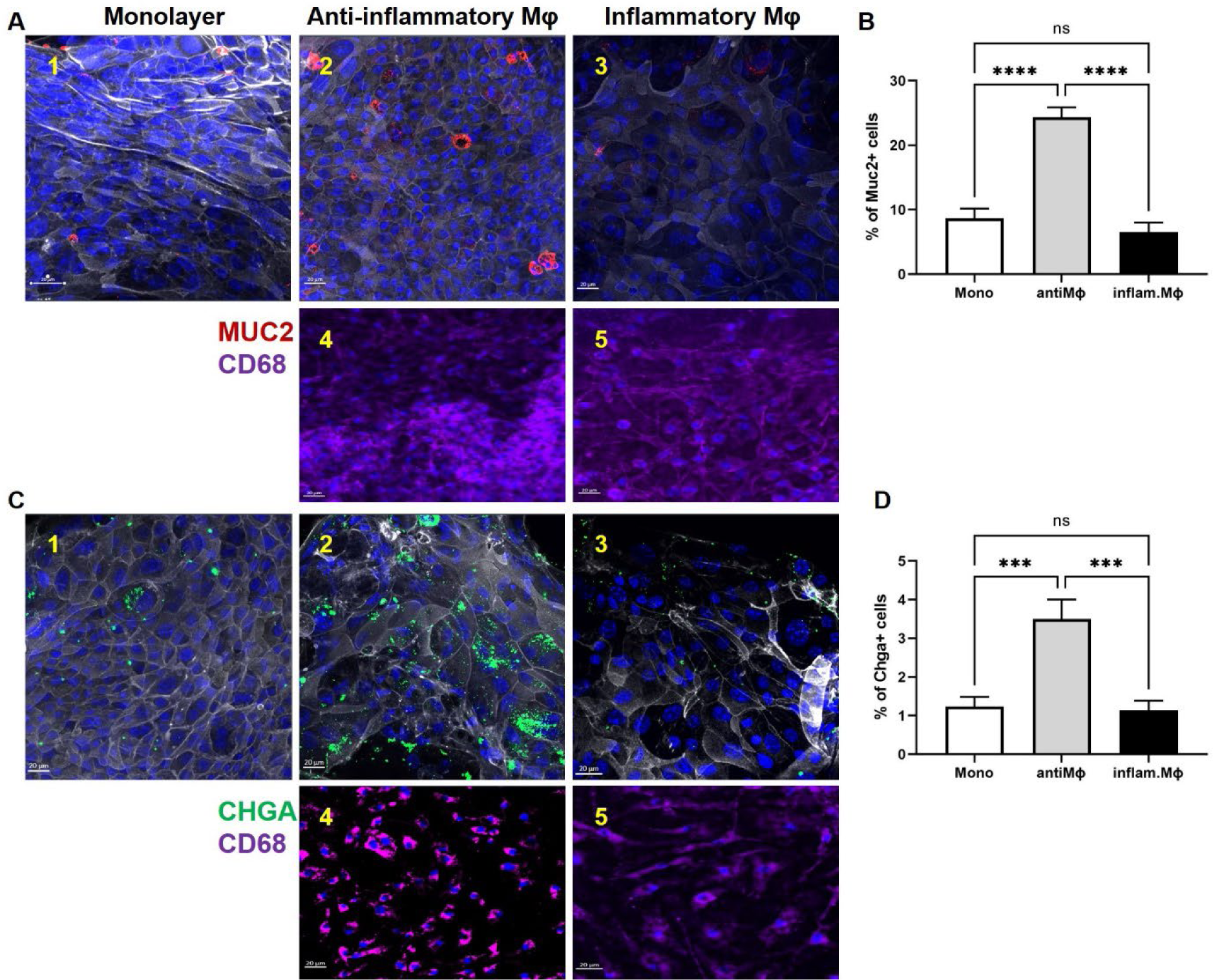
Inflammatory macrophages inhibit secretory lineage differentiation. Representative images showing mucin 2 (MUC2) and chromogranin A (CHGA) detection in mouse epithelia-macrophage (CD68) co-cultures after ISC differentiation. **(A)** MUC2 (Red) staining in 1) epithelial monolayers, 2) co-culture with anti-inflammatory macrophages (Mφ), 3) co-culture with inflammatory macrophages, 4 and 5) staining of CD68 (Purple) on the basolateral surface. **(B)** Graph shows the proportion of MUC2^+^ cells out of 1000 cells counted in each condition. **(C)** CHGA (Green) staining in 1) epithelial monolayers, 2) co-culture with anti-inflammatory macrophages (Mφ), 3) co-culture with inflammatory macrophages, 4 and 5) staining of CD68 (purple) on the basolateral surface. **D)** Graph shows the proportion of CHGA^+^ cells out of 1000 cells counted in each condition. Representative of two independent experiments, n=3 per group. Graphs indicate mean (±SD). ***P<0.001, ****P<0.0001, ns, not significant. Significance was calculated by one-way ANOVA.

Intestinal macrophages are close to the epithelial barrier, performing tissue remodeling and producing soluble factors such as prostaglandin E^2^ and Wnt ligands to maintain the integrity of the barrier^39-42^. CX3CL1-producing epithelial cells and CX3CR1-expression in intestinal macrophages mediate this tissue localization^43-45^. Other studies have utilized macrophage (U937) and colonic epithelial (Caco-2 and HT29) cell lines and shown similar results. U937 cells were skewed to inflammatory cells expressing Notch ligands and were able to turn on Notch in both Caco-2 and HT29 cell lines^46^. Although these are differentiated cell lines and cannot commit to lineage differentiation this suggests inflammatory macrophages are likely to impact the mature epithelium. Taken together, our data demonstrates anti-inflammatory (low Notch ligand expression) macrophages enhance secretory lineage differentiation while inflammatory macrophages (high Notch ligand expression) inhibit secretory lineage differentiation.

### Notch activation inhibit secretory lineage differentiation in human colonoids

Inflammatory macrophages produce proinflammatory cytokines^47^ (i.e., TNFα, IL-12, IL-23, and IL-1) that can activate inflammatory cells, increase intestinal permeability by dysregulating tight junction proteins as well as induce apoptosis in epithelial cells^48-58^. Notably, these cytokines are the current targets for biological therapy and are at the forefront of IBD treatment. Organoids derived from UC patients show defective goblet cell and mucus production and TNFα was shown to exacerbate this phenotype ^59^. In our co-culture studies, inflammatory macrophages were washed before co-culturing but we cannot rule out the effects of cytokines produced during our *in vitro* experiments. Therefore, we wanted to directly assess the role of Notch activation on goblet and enteroendocrine cell development in human colonoids. Human colonoids were allowed to differentiate for 6 days while media was replenished every 48 hrs. During this period, Notch signaling was activated in colonoids using Yhhu3792 (5-(3-Methoxyphenoxy)-N2-[4-(1-methylethyl) phenyl]-2,4-quinazo-linediamine hydrochloride)^60^ or inhibited using SAHM1, a modified peptide that disrupts protein-protein interactions and prevents notch complex assembly^61^. Using RNAscope and confocal imaging, we confirmed that Notch activation led to upregulation of the Notch target *HEY1* (**Fig. 4 A, B**) while decreasing the expression of secretory genes *MUC2* (**Fig. 4 C, D**) and *CHGA* (**Fig. 4 E, F**). Interestingly, there was no change in secretory lineage expression upon Notch inhibition when comparing unstimulated colonoids (**Fig. 4 A-F**); however, there was a significant change in secretory marker expression between Notch activation and Notch inhibition (**Fig. 4 B, D, F**). Taken together, this supports our co-culture system that Notch activation on ISC prevents secretory cell development.

**Figure 4.**
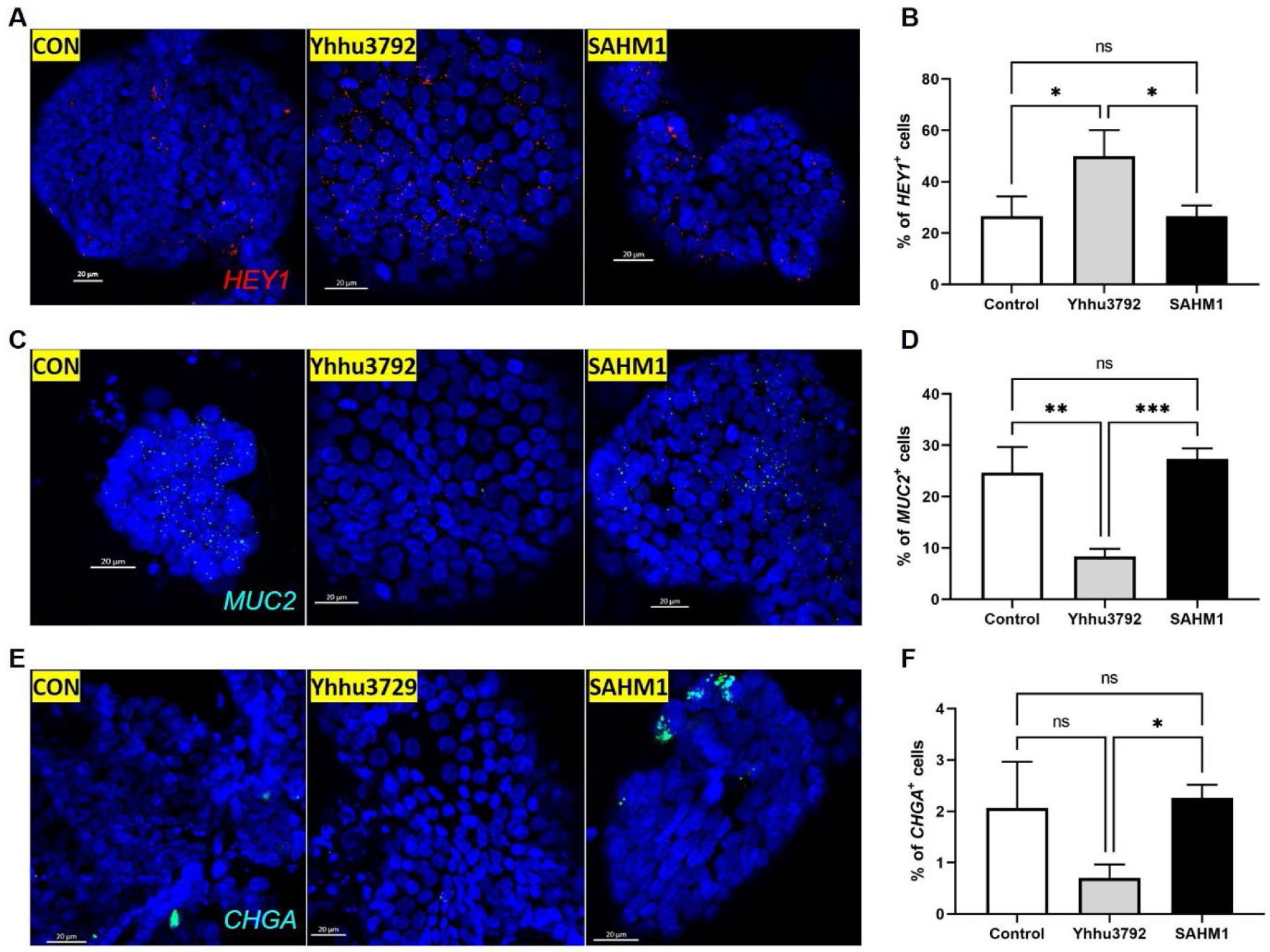
Notch activation decrease secretory lineage marker expression in human colonoids. Macrophage-free Notch activation in Human colonoids. Colonoids were treated with 10 μM of the Notch activator Yhhu3792 or 40 μM of the Notch inhibitor SAHM1 for 6 days. Notch target *HEY1* and secretory lineage markers *MUC2* and *CHGA* were quantified via RNA scope. **(A)** RNAscope detection of *HEY1* in human colonoids after left untreated or treated with Yhhu3792, or SAHM1. **(B)** Graph shows the proportion of *HEY1*^+^ cells out of 1000 cells counted in each condition. **(C)** RNAscope detection of *MUC2* in human colonoids after left untreated or treated with Yhhu3792, or SAHM1. **(D)** Graph shows the proportion of *MUC2*^+^ cells out of 1000 cells counted in each condition. **(E)** RNAscope detection of *CHGA* in human colonoids after left untreated or treated with Yhhu3792, or SAHM1. **(F)** Graph shows the proportion of *CHGA*^+^ cells out of 1000 cells counted in each condition. Representative of two independent experiments, n=3 per group. Graphs indicate mean (±SD). * P<0.05, **P<0.01, ***P<0.001, ns, not significant. Significance was calculated by one-way ANOVA.

Previous reports in IBD patient samples identified Notch activity in different compartments of the epithelium (*i.e.,* Notch activity in crypts and differentiated cells)^5,46^. The loss of Notch signaling via genetic or chemical inhibition results in intestinal inflammation and constitutive Notch signaling is found in IBD and GI cancers. Pan-Notch inhibitors being utilized for cancers are also associated with GI toxicity, specifically, goblet cell hyperplasia^1^. Nevertheless, it is unclear what component of the Notch pathway contributes to intestinal disease or what cell(s) contributes to continuous Notch activation. Here, we show in a model of intestinal inflammation and ISC differentiation, inflammatory macrophages have increased expression of Notch ligands that can prevent goblet and enteroendocrine cell differentiation (**Suppl. Fig. 5**).

Changes in enteroendocrine cell numbers as well as hormones and peptides (both increased and decreased) have been noted in UC and CD, but the mechanism(s) driving these changes are largely unknown. This study suggests Notch signaling from pro-inflammatory macrophages plays a role in enteroendocrine cell changes during intestinal inflammation. Given that additional cells such as dendritic cells and monocytes express Notch ligands we cannot rule out the influence of these cells on ISC differentiation during intestinal inflammation. Additionally, the Notch pathway also promotes proinflammatory T cell activity and can also dampen regulatory T cell function, which could drive IBD pathogenesis ^15,62-68^. The interaction of Notch ligand-positive inflammatory macrophages with T cells could also prove problematic for IBD patients. Overall, dysregulated Notch signaling appears to perturb intestinal homeostasis on multiple fronts. Future work will entail understanding the signals of anti-inflammatory macrophages impacting ISC and how inflammatory (Notch ligand-positive) macrophages affect immune cells in the gut.

## Methods

### Animals

B6.129P2-*Il10^tm1Cgn^*/J mutant (*Il10*^-/-^) (002251) and C57BL/6J (000664) mice were purchased from JAX. Both female and male mice between 14-18 weeks of age were used for experiments. All experiments were approved by the Institutional Animal Care and Use Committee of the University of New Mexico Health Sciences Center (IACUC # 20-201025-HSC), in accordance with the National Institutes of Health guidelines for use of live animals. The University of New Mexico Health Sciences Center is accredited by the American Association for Accreditation of Laboratory Animal Care.

### Macrophage-Epithelial Co-culture

Crypt isolation and colonoid generation was done as previously reported^69^ utilizing proximal colon tissue from B6 mice. Briefly, colonic tissue was minced into 0.5 cm pieces and washed several times in cold freshly prepared complete chelating solution (CCS; 5.6mM Na2HPO4, 96.2mM NaCl, 1.6mM KCl, 8mM KH2PO4, 43.4mM sucrose, 0.5mM DL-dithiothreitol and 54.9mM D-sorbitol). Colonic tissue was incubated for one hour in 10 mM EDTA in CCS on an orbital shaker at 4°C. Supernatant containing crypts were collected and centrifuged at 1500rpm for 10 minutes at 4°C. Crypts were resuspended in Matrigel (Corning, Tewksbury, MA) and plated in 24-well plates (Corning) as 30 μl droplets or utilized for flow cytometry. After polymerization at 37°C, 500 μl of expansion media (EM), described in supplemental material) was added to each well. Media was replenished every 48 hours. All colonoid cultures were maintained at 37°C with 5% CO^2^ and passaged every 5 days. To seed Transwells (0.4 µM pore size), colonoids were harvested from Matrigel using Cultrex Organoid Harvesting Solution as previously described^70^. Colonoids were resuspended in 500 μl of 0.25% trypsin, washed and transferred to 24 well plates. The plates were placed in a spinoculator at 37°C and rocked at 600 rpm for 45 minutes to digest the colonoids into single cells. Digested colonoids were transferred into 15ml conical tubes, containing 10ml PBS and spun down to pellet the cells. The supernatant was removed and single cells were resuspended in 100 μl of filter media and transferred onto collagen-coated Transwells (described in supplemental material). Seeded Transwells were placed in 24 well plates and 600 μl of filter media was added to the bottom of the plate. Both the apical and basal media were replenished every 48 hours until the monolayers were 80-90% confluent. To seed the confluent layer with macrophages, Transwells were carefully flipped utilizing sterile forceps and ensuring the media inside the Transwell did not pour out. Macrophages were generated as described in supplemental material and previously reported^26,33^ and resuspended in filter media at a density of 2000 macrophages per microliter. Macrophages suspended in filter media (50 μl) were carefully added to the bottom of the filter to cover the entire surface. The plate was carefully transferred to a cell culture incubator at 37°C for two hours. After, the Transwells were reinserted into the wells of a 24 well plate. Fresh base media (600 μl) to allow for differentiation was added to the bottom of the well while 100 μl was added into the apical side of the Transwell. The co-culture was left for 48 hours at 37°C with 5% CO^2^. The apical and basal media was removed, and the monolayer and macrophages were rinsed twice with PBS, fixed with 4% paraformaldehyde and immunostained (**Suppl. Tables S1 and S4**) for confocal microscopy.

### Human Tissue Collection and Human Colonoid Generation

Human colonoid studies were reviewed and approved by the Johns Hopkins University School of Medicine Institutional Review Board (IRB# NA_00038329) and University of New Mexico Institutional Review Board (IRB# 18-171). Colonic biopsies were obtained from healthy individuals undergoing screening colonoscopies who had given informed written consent. Colonic crypt isolation and colonoid generation were prepared as previously reported ^69,71^. Briefly, biopsy tissue was minced, washed several times in freshly prepared cold chelating solution (CCS; 5.6mM Na^2^HPO^4^, 8mM KH^2^PO^4^, 96.2mM NaCl, 1.6mM KCl, 43.4mM sucrose, 54.9mM D-sorbitol, and 0.5mM DL-dithiothreitol) and incubated 1 hour at 4°C in 10 mM EDTA in CCS on an orbital shaker. Isolated crypts were resuspended in Matrigel (Corning) and 30 μl droplets were plated in a 24-well plate (Corning). After polymerization at 37°C, 500 μl of expansion media (EM) was added for 2 days (Advanced Dulbecco’s modified Eagle medium/Ham’s F-12 (ThermoFisher, Waltham, MA), 100 U/mL penicillin/streptomycin (Quality Biological, Gaithersburg, MD), 10 mM HEPES (ThermoFisher), and 1X GlutaMAX (ThermoFisher), with 0.15 nM WNT surrogate-Fc fusion protein (ImmunoPrecise Antibodies, Fargo, ND), 15% v/v R-spondin1 conditioned medium (cell line kindly provided by Calvin Kuo, Stanford University), 10% v/v Noggin conditioned medium (cell line kindly provided by Gijs van den Brink, Tytgat Institute for Liver and Intestinal Research), 1X B27 supplement (ThermoFisher), 1 mM N-acetylcysteine (MilliporeSigma), 50 ng/mL human epidermal growth factor (ThermoFisher), 10 nM [Leu-15] gastrin (AnaSpec, Fremont, CA), 500 nM A83-01 (Tocris, Bristol, United Kingdom), 10 μM SB202190 (MilliporeSigma), 100 mg/mL primocin (InvivoGen, San Diego, CA), 10 μM CHIR99021 (Tocris), and 10 μM Y-27632 (Tocris)). After 2 days, the EM (without CHIR99021 and Y-27632) was replaced every other day. Colonoids were passaged every 10-14 days by harvesting in Cultrex Organoid Harvesting Solution (R&D Systems, Minneapolis, MN) at 4°C with shaking for 30’. Colonoids were fragmented by trituration with a P200 pipet 30-50 times, collected and diluted in Advanced DMEM/F12, centrifuged at 300 xg for 10’ at 4°C. The pellet was resuspended in Matrigel and plated as described for crypt isolation. All colonoid cultures were maintained at 37°C and 5% CO2. Unless noted, colonoid lines have been passaged >20 times.

### RNAscope

RNA was stained in colonoids via the RNAscope protocol (ACD, Newark, CA),with minor changes, as previously described^70^. Briefly, colonoids were harvested from Matrigel and fixed onto lysine coated glass slides with 4% paraformaldehyde for 1 hour. Colonoids were permeabilized with hydrogen peroxide for 10 minutes. Slides were washed 3 times with distilled water, RNAscope Protease Plus was added to submerge the colonoids. Slides were incubated for 30 minutes at 40°C in a humidified tray. Probe hybridization was done for 2 hours at 40°C. Signal amplification and detection were done as described in the RNAscope protocol. Nuclei were stained with DAPI and FluorSave Reagent (Calbiochem) was used to mount coverslips onto the slides. Images were acquired at the UNM AIM Center using the LSM 900 Zeiss AXIO Observer.Z1/7 Inverted Fluorescence Microscope with 63x oil objective lens. Images were processed using the Zen blue (version 3.4) software and ImageJ (version1.8.0) for quantification. All RNA probes used are listed in supplemental table S1.

## Statistical Analysis

Statistical analysis was performed as described in figure legends and graphs generated display mean (± SD or SEM) and were obtained using GraphPad Prism software. The data were analyzed using one-way ANOVA or two-tailed unpaired Student’s t test (Prism).

## Online supplemental material

Material and methods for colitis induction, cell and tissue preparation, organoid media, RT-qPCR, flow cytometry, ELISA, and immunostaining are found in supplemental material.

## Author contributions

RA performed all experiments and analysis with the help from ASR, AJM, SDM, JGI, and EFC. JGI participated in imaging and analysis and provided reagents for organoids. RA, ASR, and JGI participated in writing the manuscript. EFC designed the study, analyzed data and wrote the paper. All authors approved the final version of the manuscript.

## Acknowledgments

This work was supported in part by National Institutes of Health (NIH) through NIH grant no. UL1TR001449 (EFC), P20GM121176 (EFC), ES034400 (JGI), K01DK106323 (JGI) and the American Cancer Society Institutional Research Grant IRG-14-187-19 (JGI). BioRender software was used to generate graphical abstract. The data underlying this article are available in the article and in its online supplementary material.

## Supplementary Methods

### Colitis

*Il10*^-/-^ mice showed early inflammatory changes by 10-12 weeks after weaning and more severe inflammation by 14-16 weeks. Dextran Sodium Sulfate (DSS, colitis grade, ∼30,000 -50,000 MW; MP Biomedicals) was used for both acute and chronic DSS-induced colitis in C57BL/6J female mice between 8-16 weeks of age. In the acute model, mice were provided 2.5% DSS in drinking water ad-libitum for 7 days and then switched to drinking water for 7 days. In the chronic model, mice received three cycles of 2.5% DSS in drinking water ad-libitum for 7 days and each cycle was separated with two weeks of drinking water. Stool was utilized to detect lipocalin-2 (LCN-2). Stool samples were weighed and diluted in 0.1% PBS-Tween. LCN-2 ELISA was performed using Quantikine ELISA Kits (R&D Systems) according to the manufacturer’s specifications.

### Cells and tissue preparation

Colonic epithelial cells and lamina propria (LP) macrophages were isolated from the colon by a three-step process ^1-3^. 1) Separation of the colonic epithelium from the LP by chelation using 2 mM EDTA and 0.5mM dithiothreitol in HBSS buffer; 2) digestion of the remaining colon with collagenase and dispase; and 3) enrichment of CD45^+^ cells using biotinylated CD45 antibody and streptavidin beads from the digested fraction. Colonic epithelial cells were cryopreserved in RNA-Later (Invitrogen) for RNA isolation. LP macrophages were used for flow cytometric analysis. Mesenteric lymph nodes (mLN) were collected and manually homogenized into single cell suspensions. mLN cells were resuspended in RPMI prior to plating and then stimulated with phorbol 12-myristate 13-acetate (PMA, 50 ng/mL) and Ionomycin (500 ng/mL) in the presence of Brefeldin A/GolgiPlug (1:1000, BD Biosciences) and Monesin/GolgiStop, (1:1000, BD Biosciences) in preparation for intracellular cytokine staining and analysis by flow cytometry.

### Generation of primary macrophages

The generation of primary macrophages were prepared as briefly described ^3,4^, marrow was collected from the femur and tibia, washed and differentiated in DMEM (Gibco), FBS (VWR), and murine L929-fibroblast supernatant containing CSF-1 for 10 days, after which cellular morphology was evaluated to confirm differentiation. Macrophages were rested for 16 hours in media without CSF-1 prior to stimulation with LPS (200 ng/mL, Thermofisher) or IFNγ (10 ng/mL, Peprotech) overnight. Macrophages were used for flow cytometric analysis or co-culturing with epithelial cells derived from organoids. Prior to co-culturing with colonic epithelial cells, macrophages were washed twice before incubation.

### Flow Cytometry

Bone marrow-derived or LP cells were pretreated with Stain FcX (anti-CD16/32) (Biolegend) before being stained for cell surface markers. For staining, cells were washed with FACS buffer [90% by volume 1x PBS, (Life Technologies), 10%FBS (VWR), and 0.05% 0.5 M EDTA, (Invitrogen)] prior to staining. Cells were incubated for 20 min with antibody stain in FACS buffer, followed by washing with FACS buffer, and fixation in 1% PFA. To examine intracellular proteins, cells were permeabilized and incubated with antibodies targeting cytokines for 30 minutes and then washed and resuspended in FACS buffer. Sample analysis was performed on LSR Fortessa (BD Biosciences) and analyzed using FlowJo software (TreeStar). Antibodies used for flow cytometric analysis are listed in Table S2.

### RNA Isolation, Quantification, and RT-qPCR

RNA isolation was performed on cells or tissues cryopreserved in RNA-Later (Invitrogen) using the RNA Purelink Minikit (Invitrogen) according to the manufacturer’s protocols. RNA was quantified on Nanodrop2000 and all samples yielded a 260/280 of 2 ± 0.15. cDNA synthesis was performed using Oligo(dT) Primer and SSIV Reverse Transcriptase in the presence of Cloned Ribonuclease Inhibitor (all ThermoFisher). Reverse transcription reaction and RT-qPCR runs utilized Taqman MasterMix (ThermoFisher) on a viiA7 Thermal Cycler (ThermoFisher) using QuantStudio 7. RT-qPCR primers and reagents used are listed in Table S3.

### Immunostaining for Confocal Microscopy

Colonoids were harvested from Matrigel using Cultrex Organoid Harvesting Solution as previously described^5^. Colonoids were fixed for 30 minutes in 4% paraformaldehyde (Electron Microscopy Sciences, Hatfield, PA). Colonoids were washed twice using 500 μl of 1X PBS. Colonoids were permeabilized and blocked simultaneously for 1 hour using a 10% Fetal Bovine Serum (Atlanta Biologicals, Flowery Branch, GA), 0.1% saponin (MilliporeSigma) solution prepared in PBS. After permeabilization, colonoids were washed three times in 1X PBS. 100 μl of primary antibody prepared at 1:50 dilution in 1X PBS was added to organoids and incubated overnight at 4°C. Organoids were washed 3 times with 1X PBS, and 100 μl of AlexaFluor secondary antibodies including AlexaFluor-647 phalloidin, diluted 1:200 in 1X PBS, were added for 1h at room temperature. Hoechst 33342 (ThermoFisher) 1 mg/ml was added for 5 minutes. After three 1X PBS washes, FluorSave Reagent (Calbiochem) was added to the colonoids and mounted on glass slides. Confocal imaging was performed at the UNM AIM Center using the LSM 900 Zeiss AXIO Observer Z1/7 Inverted Fluorescence Microscope. Images were processed using the Zen blue (version 3.4) software. Images were processed using the Zen blue (version 3.4). Primary and secondary antibodies used are listed on Table S4.

### Organoid Growth media

Intestinal organoid media for monolayer plating (referred to as ‘filter media’) was comprised of Advanced Dulbecco’s modified Eagle medium/Ham’s F-12 (ThermoFisher, Waltham, MA), 100 U/mL penicillin/streptomycin (Quality Biological, Gaithersburg, MD), 50% v/v WNT3A conditioned medium (ATCC CRL-2647), 15% v/v R-spondin1 conditioned medium (cell line kindly provided by Calvin Kuo, Stanford University), 10% v/v Noggin conditioned medium (cell line kindly provided by Gijs van den Brink, Tytgat Institute for Liver and Intestinal Research), 1X B27 supplement (ThermoFisher), 10 mM HEPES (ThermoFisher), 1X GlutaMAX (ThermoFisher), 1mM N-acetylcysteine (MilliporeSigma), 50 ng/mL human epidermal growth factor (ThermoFisher), 10 nM [Leu-15] gastrin (AnaSpec, Fremont, CA), 500 nM A83-01 (Tocris, Bristol, United Kingdom), 10 μM SB202190 (MilliporeSigma), 100 mg/mL primocin (InvivoGen, San Diego, CA). Intestinal organoid media for expansion of 3-dimensional organoids (referred to as expansion media) had the same composition as filter media except for the replacement of 50% v/v WNT3A conditioned medium with 0.15 nM WNT surrogate-Fc fusion protein (ImmunoPrecise Antibodies, Fargo, ND). The volume of the conditional WNT3A was made up for by Advanced Dulbecco’s modified Eagle medium/Ham’s F-12 (ThermoFisher). Base media, for differentiation of organoids, had the same composition of filter media but lacked WNT3A, Rspo-1 and SB202190. 0.4 µM pore sizeTranswell filters (Corning) were coated using 100 µl of 100 µg rat collagen IV (Corning) at 4°C overnight, aspirated, and washed (3x) with 200 μl of sterile water prior to use.

## Supplemental Tables

**Table S1.** RNAScope probes.

| Probe | Source | Identifier |
| --- | --- | --- |
| Hs-CHGA-C4 | ACD | Cat# 311111-C4 |
| Hs-HEY1 | ACD | Cat# 311201 |
| Hs-MUC2-c2 | ACD | Cat# 3122871-C2 |

**Table S2.**
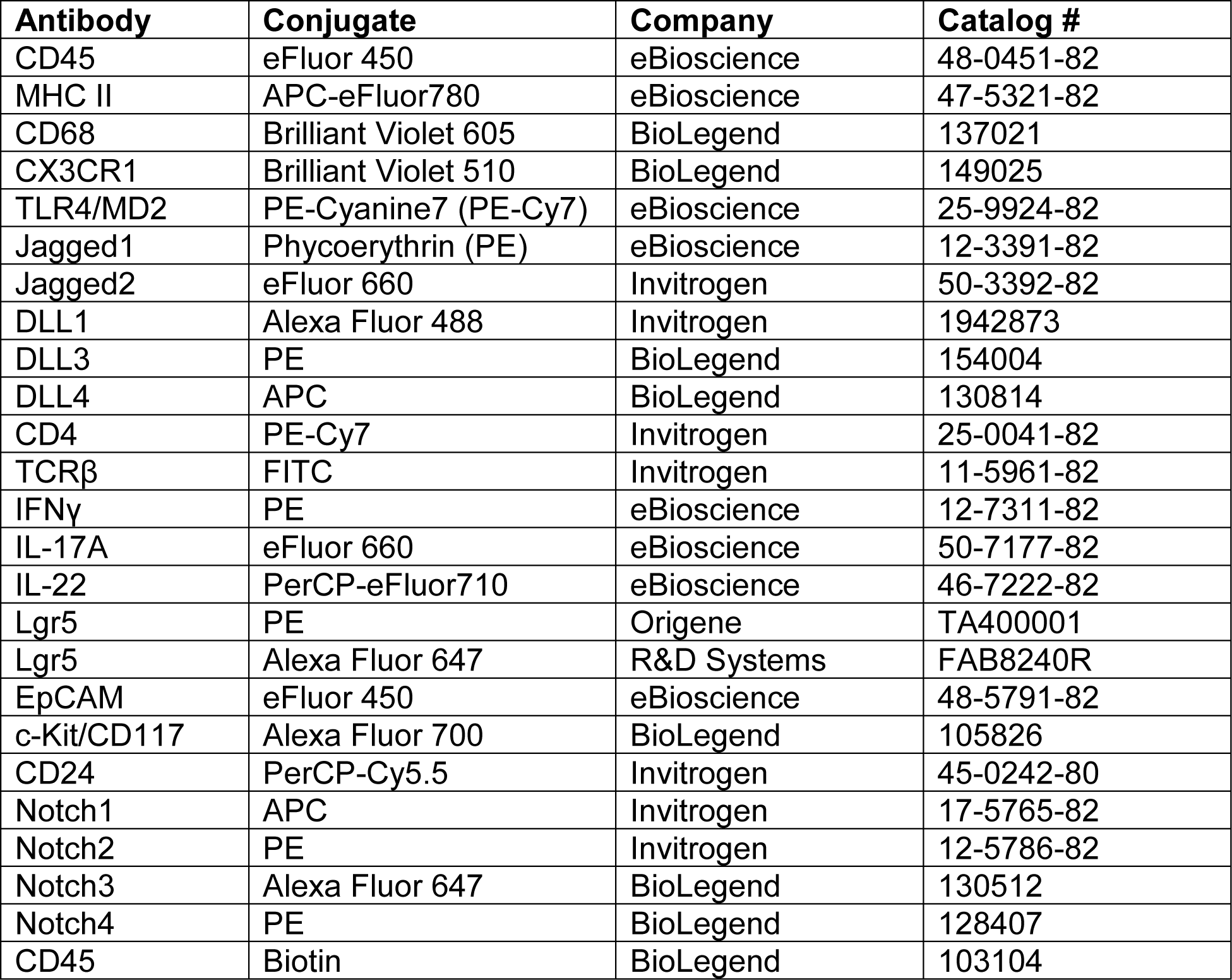
Flow Cytometry antibodies.

**Table S3.** PCR Primers.

| **Assay ID** | **Source** | **Identifier** |
| --- | --- | --- |
| Atoh1: Mm00476035_s1 | ThermoFisher | Cat# 4331182 |
| Gapdh: Mm99999915_ | ThermoFisher | Cat# 4331182 |
| Hes1: Mm01342805_m1 | ThermoFisher | Cat# 4331182 |
| Hey1: Mm00468865_m1 | ThermoFisher | Cat# 4453320 |
| Hes5: Mm00439311_g1 | ThermoFisher | Cat# 4331182 |
| Dtx1: Mm00492297_m1 | ThermoFisher | Cat# 4453320 |
| Gli5: Mm00494654_m1 | ThermoFisher | Cat# 4331182 |

**Table S4.** Primary and secondary antibodies for Immunostaining.

| **Antibody** | **Source** | **Identifier** |
| --- | --- | --- |
| CD68 | ThermoFisher | Cat# 140681-82 |
| MUC2 | Novus | Cat# NBP1 31231 |
| Chromogranin A | Santa Cruz | Cat# Sc393941 |
| Donkey anti-Rabbit IgG-AF488 | ThermoFisher | Cat# A-21206 |
| Donkey anti-Mouse IgG, AF568 | ThermoFisher | Cat# A10037 |
| Donkey anti-Mouse IgG, AF488 | ThermoFisher | Cat# A21202 |
| Goat anti-Rabbit IgG AF568 | ThermoFisher | Cat# A1101042 |
| Phalloidin-Alexa Fluor 647 | AAT Bioquest | Cat# 1000854 |

**Supplemental Figure 1.**
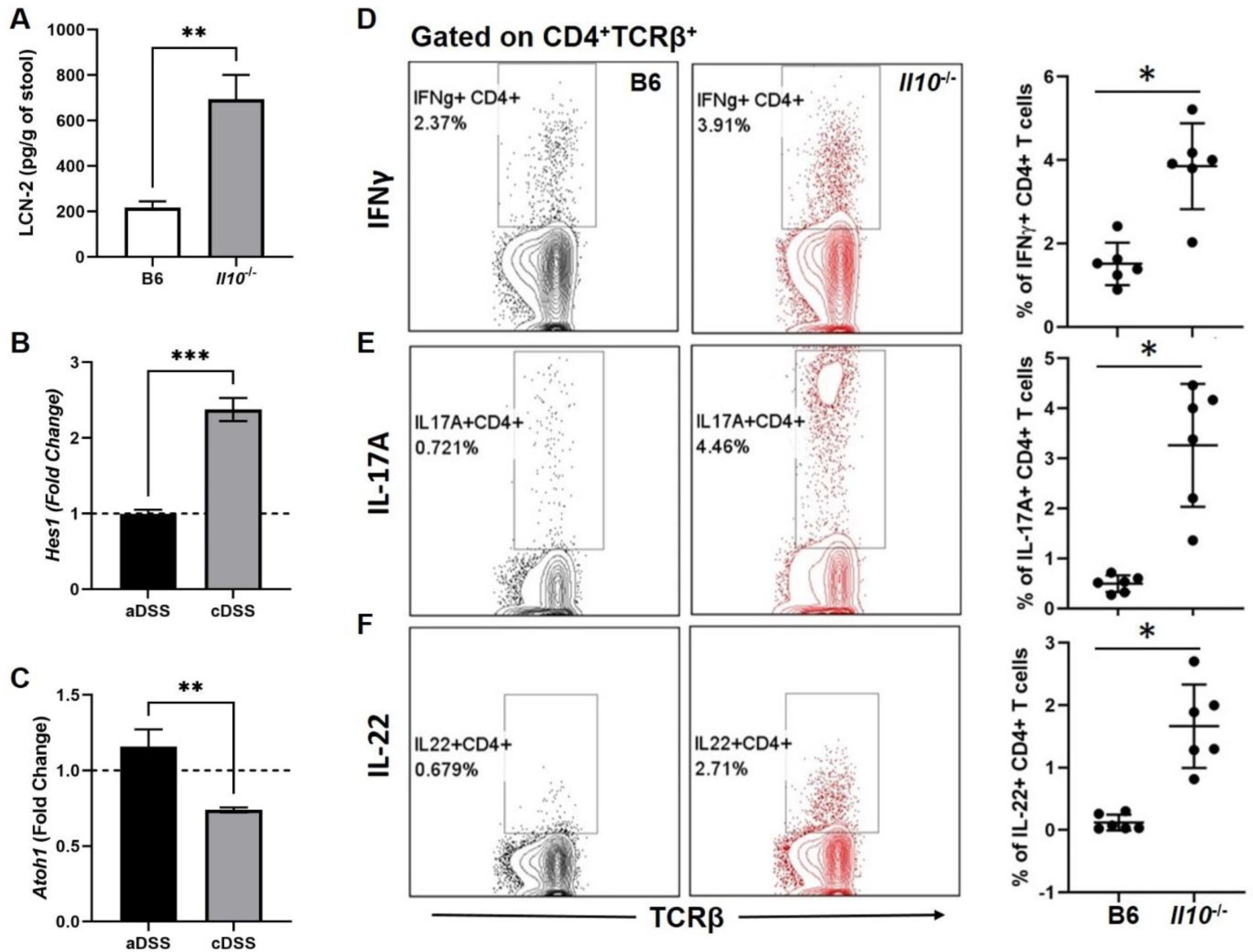
Comparison of colitic *Il10*^-/-^ and non-colitic B6 mice. **(A)** Fecal LCN-2 levels as detected by ELISA from colitic *Il10*^-/-^ and non-colitic B6 mice. Colon epithelial **(B)** *Hes1* and **(C)** *Atoh1* gene expression after acute (aDSS) and chronic (cDSS) DSS-induced colitis. **(D)** Representative dot plots showing Intracellular IFNγ staining in CD4^+^ T cells isolated from mLN in colitic *Il10*^-/-^ and non-colitic B6 mice and corresponding graph. **(E)** Representative dot plots showing Intracellular IL-17A staining in CD4^+^ T cells isolated from mLN in colitic *Il10*^-/-^ and non-colitic B6 mice and corresponding graph. **(F)** Representative dot plots showing Intracellular IL-22 staining in CD4^+^ T cells isolated from mLN in colitic *Il10*^-/-^ and non-colitic B6 mice and corresponding graph. Representative of two independent experiments, Graphs indicate mean (±SEM). *P<0.05, **P<0.01, ***P<0.001, Significance was calculated by two-tailed unpaired Student’s t test.

**Supplemental Figure 2.**
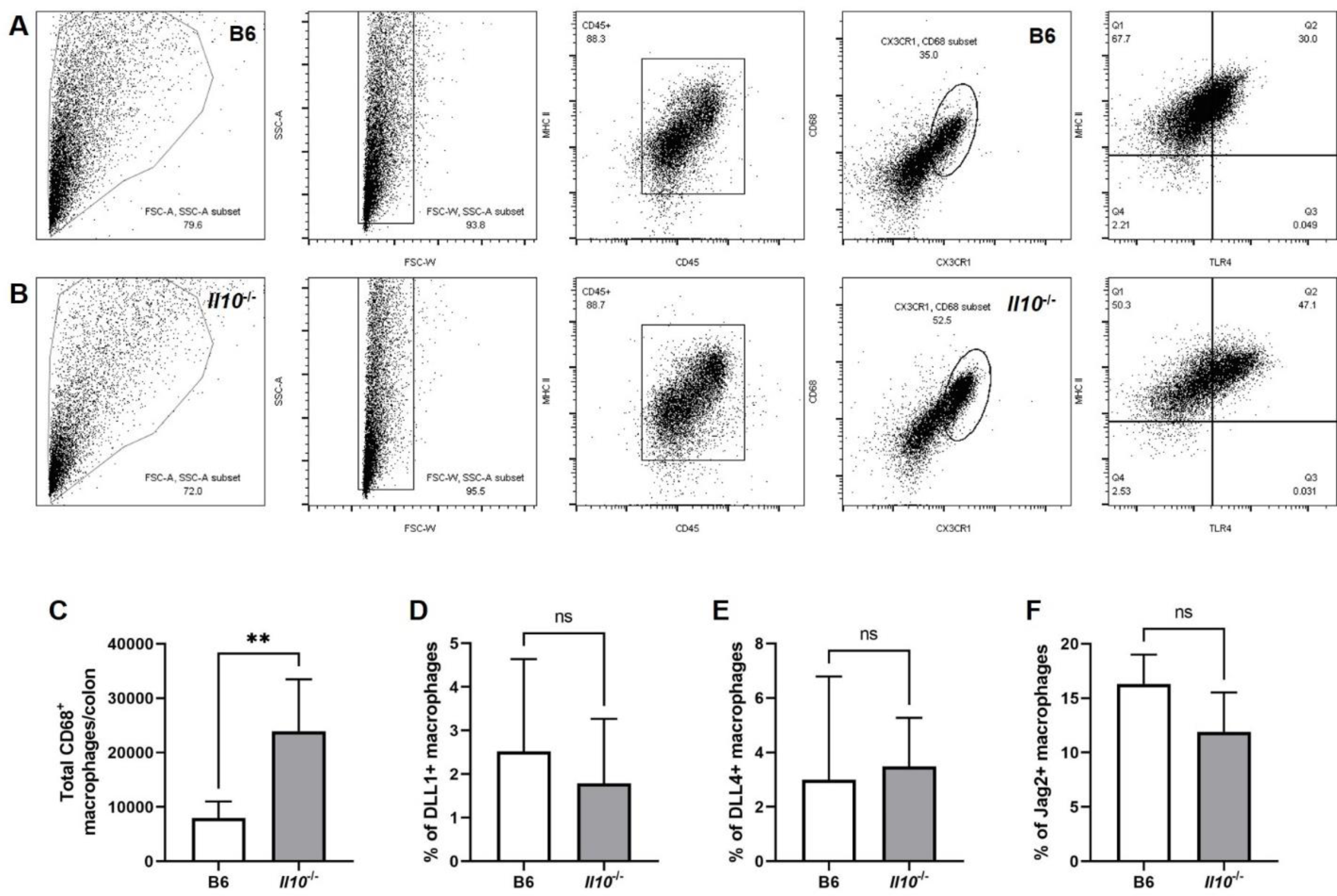
Colonic lamina propria macrophages and Notch ligand expression. Gating strategy of LP macrophages from **(A)** B6 and **(B)** *Il10*^-/-^ mice. **(A, B)** Dot plots showing FSC/SSC. FSC-W and lamina propria (gated on CD45^+^ and MHC II^+^) cells and further gated on CD68 and CX3CR1. Dot plot showing gated CD68+ cells TLR4 expression. **(C)** Graph showing total number of macrophages (MHC-II^+^CD68^+^CX3CR1^+^). **(D-F)** Graphs showing the percent of **(D)** DLL1^+^, **(E)** DLL4^+^, and **(F)** Jagged2^+^ (Jag2) macrophages. Representative of two independent experiments (n=3 per group), Graphs indicate mean (±SD). **P<0.01, ns, not significant. Significance was calculated by two-tailed unpaired Student’s t test.

**Supplemental Figure 3.**
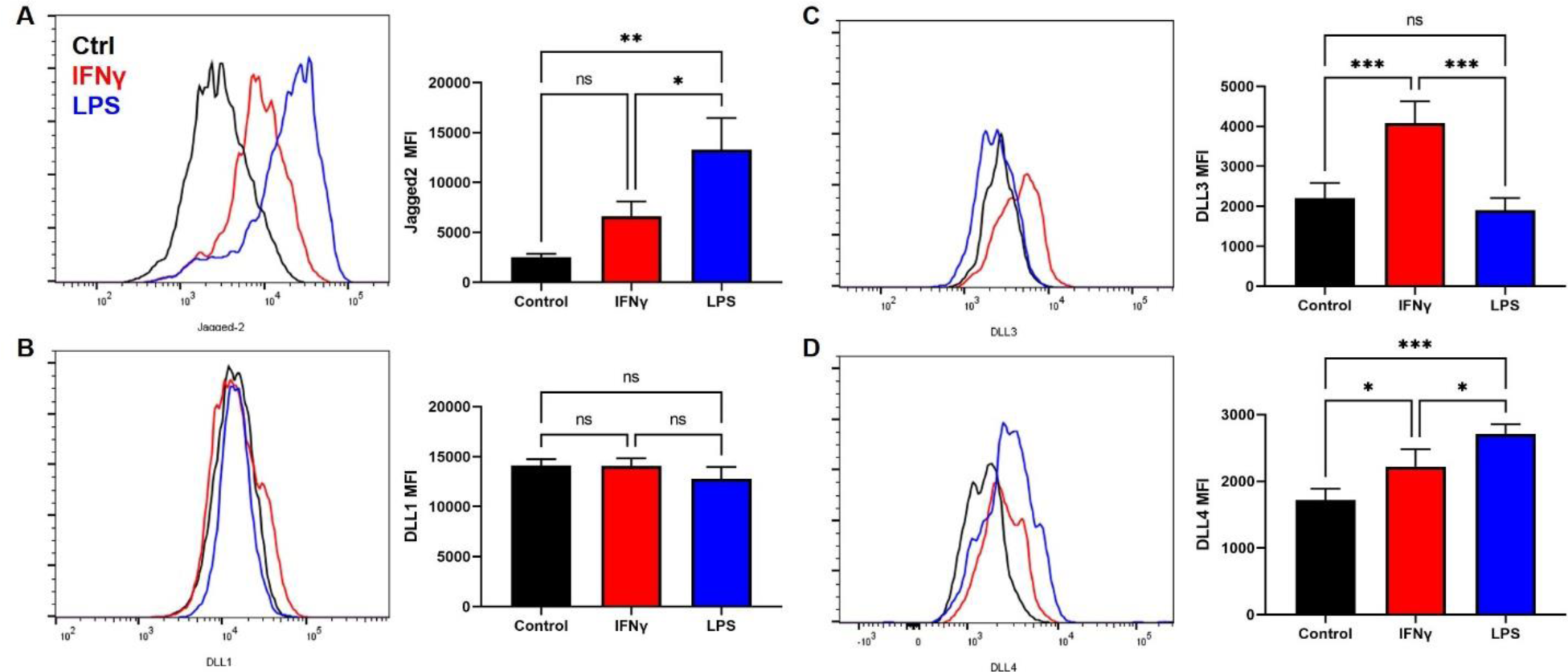
Notch ligand expression in macrophages after inflammatory stimulation. Representative histograms showing **(A)** Jagged2, **(B)** DLL1, DLL3, and **(D)** DLL4 expression in B6 macrophages stimulated with LPS (Blue) or IFNγ (Red) compared to baseline/unstimulated (Black) macrophages and corresponding graph showing the geometric mean fluorescence intensity (MFI) of **(A)** Jagged2, **(B)** DLL1, **(C)** DLL3, and **(D)** DLL4 in unstimulated and stimulated B6 macrophages. Representative of two independent experiments (n=3 per group), Graphs indicate mean (±SD). *P<0.05, **P<0.01, ***P<0.001, ns, not significant. Significance was calculated by one-way ANOVA.

**Supplemental Figure 4.**
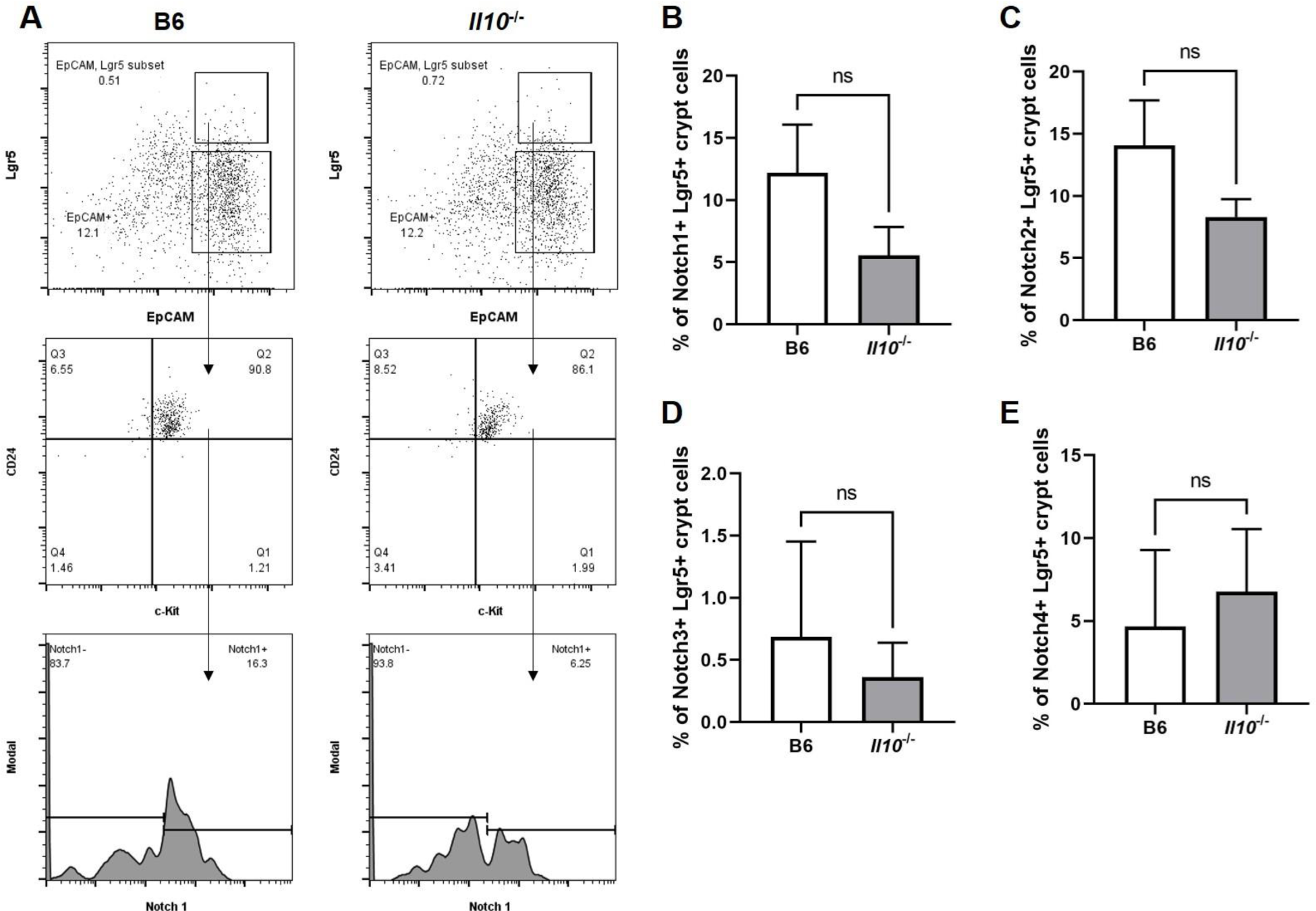
Notch Receptor on colonic crypt cells. Crypt cells were isolated from the colon of B6 or *Il10*^-/-^ mice to examine Notch receptor expression. **(A)** Gating strategy of Lgr5^+^ crypt cells – Lgr5^+^ EpCAM^+^CD24^+^c-Kit^+^ cells were examined for Notch receptor expression. Graphs showing the percent of **(B)** Notch1, **(C)** Notch2, Notch3, and **(E)** Notch4 Lgr5^+^ crypt cells isolated from B6 or *Il10*^-/-^ mice. Representative of two independent experiments (n=3 per group), Graphs indicate mean (±SD). ns, not significant. Significance was calculated by two-tailed unpaired Student’s t test.

**Supplemental Figure 5.**
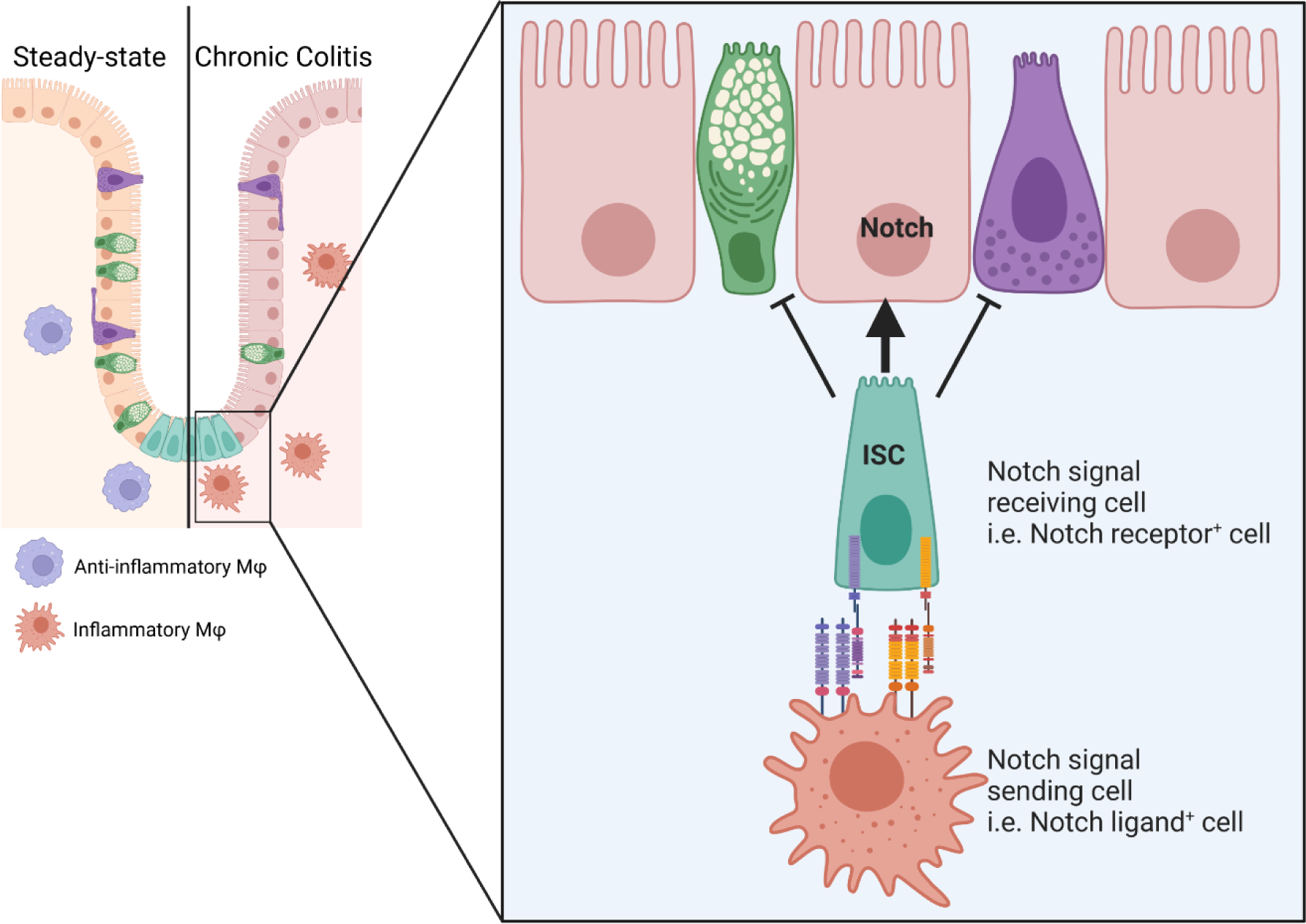
Graphical image showing inflammatory macrophages express increased levels of Notch ligands that can influence intestinal stem cells (ISC) to become absorptive cells through the promotion of Notch signaling (Image made with BioRender).

